# A validated set of neural gene reporter mice and chemical tracer tools for mapping knee innervating neurons

**DOI:** 10.1101/2025.09.29.679397

**Authors:** Ibdanelo Cortez, Carolina Leynes, Vanessa Belizaire, RE-JOIN Consortium, Nele A. Haelterman, Brendan H. Lee, Russell S. Ray

**Author notes:** Corresponding author To whom correspondence should be addressed: Russell S. Ray, Ph.D. Baylor College of Medicine Houston, TX 77030.

## Abstract

Joint pain is an increasing concern for our aging population, as current therapies to slow joint disease progression or reduce pain are largely ineffective and often carry significant health and dependency risks. Age and joint disease induce changes to all tissues that make up the joint, including the dense neural network that innervates the joint. Several studies have correlated joint innervation changes in diseases such as osteoarthritis or rheumatoid arthritis, but little is known about their respective functional consequences. How subtypes of knee-innervating neurons affect pain experience remains relatively uncharacterized. A few studies focused on a single neural subtype due to the limited availability of validated tools to study joint innervation. To better understand the relationship between aging, joint disease, and pain, systematic characterization of nociceptors and other neural subtypes regulating joint homeostasis and pain is urgently needed. This study’s objective was to establish a validated molecular and genetic toolbox for accurate mapping of the neuro-architecture in the murine knee. We screened genetic reporter mice, containing different combinations of Cre and Flp recombinase alleles along with recombinase responsive reporter alleles to either highlight peripheral nociceptors or post-ganglionic sympathetic neurons for their specificity and accuracy in labeling specific neural subtypes in the dorsal root ganglia, sympathetic ganglia, and the knee joint. Additionally, we compared the performance of a series of conventional retrograde tracers for effective labelling of sensory and sympathetic neurons innervating the knee joint. The validated molecular and genetic tools identified in this study will facilitate the creation of comprehensive joint innervation maps in physiological and pathological contexts, setting the stage for identifying the cellular and molecular changes responsible for mediating joint pain, a necessary goal for improved therapeutic interventions.

**Lay summary:** Joint diseases, such as Rheumatoid Arthritis and Osteoarthritis significantly affect the neural landscape in joints, impacting pain, balance, and joint health. Understanding these nerve changes can provide insights into the drivers of joint pain and potential treatments. Our study evaluated genetic and conventional neural tracer tools to visualize and track knee innervating nerve fibers. We found two genetic mouse lines that specifically highlight sensory and sympathetic nerves, making them suitable models for knee joint studies. Additionally, the fluorescent tracer True Blue, effectively marks cell bodies of knee-innervating neurons. These tools will help researchers better understand nerve changes in painful joint pathologies.

## Introduction

The incidence of chronic musculoskeletal pain in patients with joint diseases, such as osteoarthritis (OA), low back pain, rheumatoid arthritis, and others, continues to rise, underscoring a critical need for our fundamental understanding of the neural networks and molecular mechanisms that mediate pain signals from joint tissues.^1,2^ OA is a leading musculoskeletal condition that can affect multiple joints, contributing to years lived with disability^2,3^. Symptoms such as swelling, stiffness, and pain are typically managed with broad-spectrum therapies that provide only temporary relief and fail to halt joint degeneration^4,5^. As disease progresses and treatments remain inadequate, a growing number of treatment-resistant patients present with chronic pain, prolonged disability or need for surgical intervention^6^. As pain becomes refractory to first-line treatments, caregivers often prescribe opiates.. To address and minimize the use of opiates in treating osteoarthritic pain and other joint pain, the National Institutes of Health has developed the RE-JOIN program. RE-JOIN is a consortium of 25 + affiliated groups, whose goal is to map the neural architecture of the healthy and diseased knee and mandibular joint to clear path of improved therapeutic efficacy and pain treatment for patients with OA.

Our current understanding of the neuronal networks that innervate the knee is primarily based on anatomical and histological studies, identifying neuronal subtypes by labeling peptide expression and myelination. These studies revealed sensory and sympathetic neural innervation of several joint tissues, including the synovium, infrapatellar fat pad, menisci, and subchondral bone^7^. Nociceptive neurons, the sensory neurons relaying pain signals, are characterized by morphological features such as cell size and myelination status, as well as the production of nociceptor sensitizing peptides (peptidergic versus non-peptidergic)^8,9,10,11^. For example, Delta A fiber type nociceptors are medium-small, myelinated neurons that relay acute pain signals. Conversely, C-fiber type neurons are unmyelinated-small neurons that transmit chronic dull/achy types of pain^12,13^. Sympathetic fibers are primarily involved in peripheral vasoactivity and are identified by colocalization of tyrosine hydroxylase with vascoactive intestinal peptide (VIP) vasodilating neurons or with Neuropeptide-Y vasoconstricting neurons (NPY)^14,15^. Importantly, several groups have shown that knee innervation changes in human and animal models of OA, with ectopic sprouting occurring in several joint tissues including the synovium and subchondral bone^13,16–20^. In addition, single-cell RNA sequencing (scRNA-seq) of lumbar DRGs revealed OA-induced changes in subpopulations of peptidergic and nonpeptidergic neurons based on gene expression. These neurons are thought to mediate nociceptive signaling, but other populations and mechanisms that may contribute directly or indirectly to ongoing pathogenesis and pain remain largely understudied^7^. For example, distinct peptidergic and non-peptidergic nociceptive neurons undergo physiological and architectural changes in the OA knee, but the functional consequences of these changes, or their correlation with pain experiences, are not fully understood.

In parallel, some studies have shown that sympathetic innervation of the knee similarly responds to OA-induced alterations in joint homeostasis, and sympathetic neurons are thought to modulate sensory neural function to indirectly modulate pain in chronic pain conditions^21,22^. While these initial studies have provided a window into the vast innervation patterns that permeate the joint, the use of immunohistochemistry and antibodies for profiling neural subtypes, limits the anatomical resolution, opportunities for further molecular profiling, and functional interventions for building multifaceted anatomical, genetic, molecular, and functional maps in the healthy and diseased joints^18,23^.

Conditional genetic reporter models have proven to be a powerful approach to visualize and further characterize the cell specific neuroarchitecture in adult mice^24–26^. For example, mouse genetic fluorescent reporter systems have transformed our ability to visualize neural networks by providing robust, stable labeling of neuronal somata, dendrites, and nerve terminals across peripheral and central nervous systems^26^. By placing fluorescent reporters directly or indirectly (i.e. using recombinases) under the control of neural-subtype–specific promoters, researchers can trace distinct circuits within peripheral tissues. Furthermore, genetically distinct neural populations are now accessible with intersectional genetic techniques, allowing interrogation of novel subsets of neurons in a diverse and widespread population^24,25^. In addition to anatomical studies, such genetically delineated neuronal populations can be unambiguously characterized in terms of their electrophysiology, molecular profiles (i.e. transcript- and prote-omics), and with the use of opto- and chemogentics, functionally interrogated.

The knee joint, however, is structurally complex and is innervated by a heterogeneous array of neurons that govern sensation, proprioception, and tissue homeostasis^27^. Capturing the full neural landscape of the knee will require extensive characterization of several genetic reporter lines to improve upon traditional nociceptive and tyrosine-hydroxylase (i.e. sympathetic) immunostaining techniques. As powerful as genetic approaches are, potential off-target labeling from random transgene insertion, non-specific tissue promoter activity, or germline Cre recombination are possible risks for cell lineage tracing and thus require thorough characterization prior to studying changes in physiological or pathological contexts. Genetically engineered mouse lines expressing Cre recombinase in peripheral neurons are invaluable tools to provide insights into the neuroarchitecture and function in tissues. Several transgenic mouse lines are currently available to study peripheral neuron populations using pan-neuronal proteins or study subpopulations based on nociception and somatosensation. Pan-neuronal recombinase mouse driver lines like *Avil^tm^*^2^(cre)*^Fawa/J^*(Advillin-Cre) combined with recombinase responsive reporters mouse lines can be very useful for visualizing peripheral neurons in tissues and regions such as the knee joint where neural characterizations using immunohistochemistry are difficult^28^. To study somatosensory populations, *Trpv1^tm^*^1^(cre)*^Bbm/J^*(Trpv1-Cre) and *Piezo2^tm^*^1.1^(cre)*^Apat/J^* (Piezo2-Cre) driver mouse lines are commonly used for studying thermal sensation, mechanosensation and nociception signaling^29,30^. Furthermore, the *Scn10a^tm^*^2^(cre)*^Jwo/TjpJ^* (Scn10a-Cre)-(NaV1.8) mouse line provides critical insight for pain biology and drug studies^31^. The *E2f1^Tg^*(Wnt1–cre)*^2Sor/J^* (Wnt1-Cre) mouse line is widely used to study fate-mapping of neural crest derived cells^32–34^. Sympathetic innervation in the knee regulates several processes that maintain joint homeostasis, yet mapping this population is typically done with tyrosine hydroxylase (TH) immunostaining, which also labels a subset of sensory neurons. Here, we characterized two noradrenergic driver lines, the *Tg(Dbh-cre)^KH^*^212^*^Gsat/Mmucd^* (DBH-Cre) and *Dbh^em^*^1.1^(flpo)*^Rray^* (DBH-p2A-Flpo) to genetically label noradrenergic neural patterns. Despite their use in the literature, information on their neural specificity, particularly in the knee, remains incomplete.

Retrograde neural tracing offers a third effective strategy for mapping neuronal networks, particularly for defining the projection targets of neurons of interest; this approach has been successfully applied across a range of tissues^35^. Indeed, a wide array of these chemical dyes has been validated for neuroanatomical studies in the peripheral and central nervous systems (PNS; CNS); most reach cell bodies quickly, preserve neuronal viability, and provide high-contrast labeling of neuronal somata. Similar conventional retrograde tracing approaches have been applied to study knee joint innervation, typically involving intra-articular injection of the selected dye followed by imaging of backlabeled neuronal cell bodies in the dorsal root ganglion (DRG) ^5,17,14,36,37,38^ However, multiple types of conventional tracers exist, with distinct molecular profiles and uptake. For example, stilbene derivatives Fast Blue, True Blue and Fluorogold are commonly used due to their low toxicity, and effective somal labeling of afferent projections. The protein/toxin fluorophore derivative Cholera Toxin B-Alexa Fluor is a receptor mediated endocytosed tracer commonly used due to quick transport and long distance tracing. Microsphere latex Retrobeads^®^ are becoming more popular for their low toxicitiy, permanent labeling and minimal diffusion at injection sites^40^. Although these dyes have been validated in the CNS and skeletal muscle, a comprehensive study comparing these tracers and their ability to label sensory and sympathetic neurons innervating joint tissues has thus far not been reported^40,41^. A validated musculoskeletal neural mapping toolkit, consisting of robust, effective intra-articular retrograde tracers and characterized genetic reporter lines that specifically label neuronal subtypes, would benefit the field by providing a set of tools that, together, clear a path toward high-resolution neuroanatomical mapping, molecular profiling, and functional interrogation. To build such a toolkit, the present study evaluated commercially available genetic strains expressing cell specific Cre and FLP recombinases, combined with their respective genetic reporters to label sensory and sympathetic neuronal subtypes. Further, popular conventional retrograde tracers were evaluated for their ability to label specific populations innervating the knee joint. Importantly, these validated tools can be combined using intersectional genetic approaches for even greater genetic resolution and layered with retrograde tracing to yield high-resolution genetic and spatial tracing results and producing comprehensive, genetically defined projection maps of knee innervation across physiological and pathological conditions, setting the stage for future experiments that will clear a path for key mechanistic insights into pain development in OA and other joint diseases.

## Materials and Methods

### Animals

Male and female mice in this study were cared for in compliance with policies administered by Public Health Service Policy on Humane Care and Use of Laboratory (PHS) and approved by Institutional Animal Care and use Committee (IACUC) at Baylor College of Medicine (protocol AN-6171). Mice were allowed free access to water and rodent chow in filter top cages that were supplemented with bedding and nesting material. Mice were housed in a 12-hour light/dark cycle (6am lights on - 6pm lights off) room that was maintained at 22°C in 30% humidity room throughout study. See table 1 for mouse line information. Subjects in this study were produced from recombinase driver and recombinase-responsive reporter strain breeders that were purchased from Jackson Laboratory or previously developed in the Ray lab. To identify robust genetic tools and resources that can be used to study sensory innervation of the mouse knee joint, we first performed a literature review and selected 7 mouse strains that had previously been reported to express Cre/Flp recombinase in neuronal subtypes (**Table 1)**. To confirm genotypes of genetic reporter mice used in this study, PCR and gel electrophoresis were performed using primers for Cre, Flp, and ROSA genes, prior to collecting tissues. For each genotype, a total of three mice were used in this study to characterize neuronal expression in knee and ganglia tissues. An exception to this was the Advillin-cre reporter line where only one mouse was used for knee analysis. This mouse line was discontinued prior to final data collection of this study due to poor breeding rates. To validate previously reported neuronal expression patterns in these lines, we crossed the selected Cre- or Flp-recombinase strains to fluorescent reporter _mice *R26*_*_CAG-LoxSTOPLox-TdTomato_* _(Ai9) or *R26*_*_CAG-FrtSTOPFrt-TdTomato_* _(Ai65F). We first assessed_ Cre- or Flp-reporter activity in the synovial, menisci and bone tissues of the knee joint, as this tissue is densely innervated.

**Table 1.**
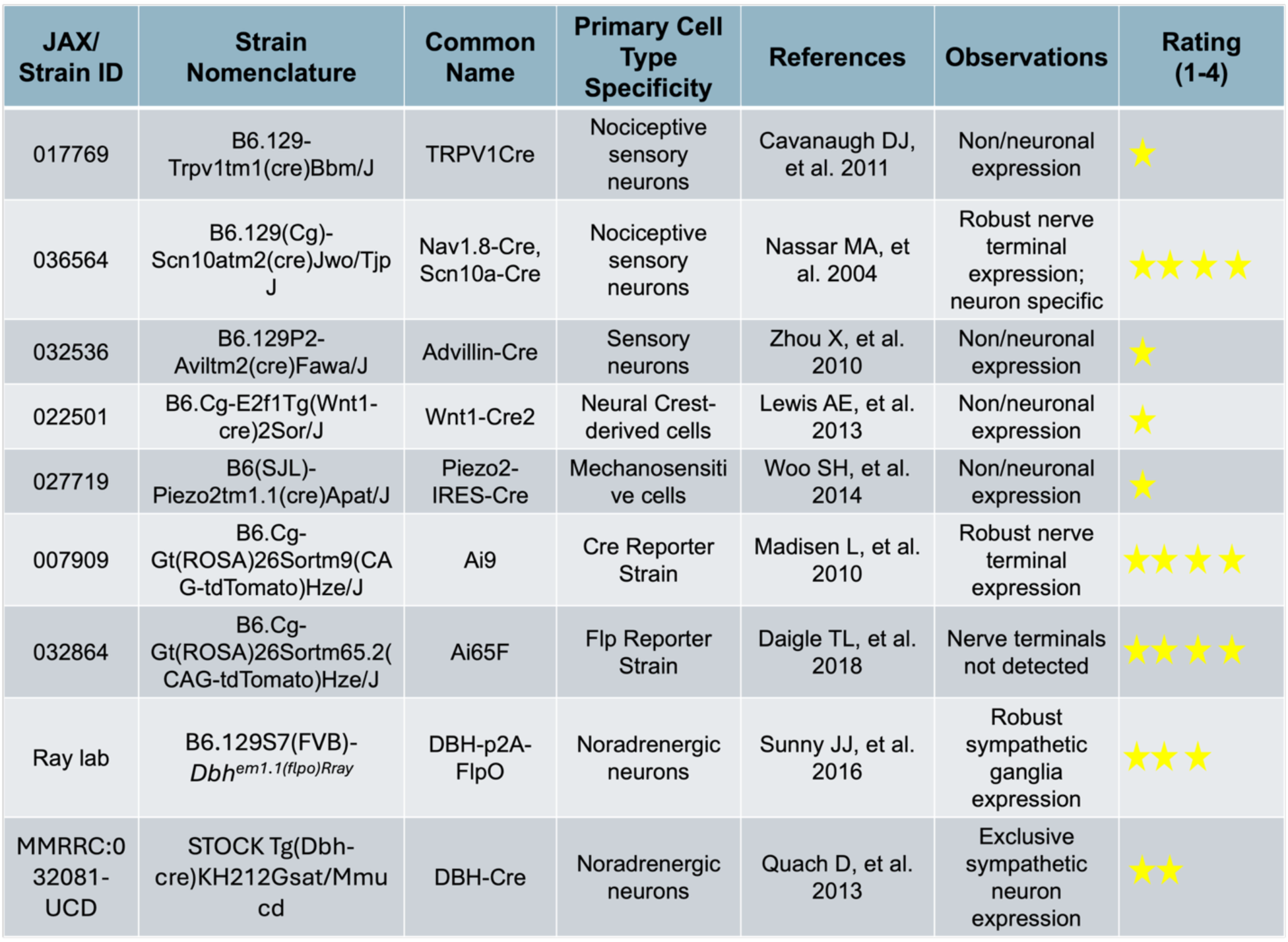

### Conventional retrograde tracers

Fast Blue (PolySciences Inc. Warrington, PA [Cat.17740]) was diluted to 1% in sterile saline and stored at -20°C. Cholera Toxin B - Alexa Fluor 647 (ThermoFisher. Waltham, MA [Cat. C34778]) was diluted to 0.2% in sterile saline and stored at -20°C. Fluoro-gold (Fluorochrome. Denver, CO [Cat. Fluor-Gold 20mg]) was diluted to 4% in sterile saline and stored at 4°C. True Blue (MedChem Express. Monmouth Junction, NJ[Cat. HY-D1161]) was diluted to 2% in 100% sterile pharmaceutical grade, DMSO (Millipore-Sigma. Darmstadt, Germany) and stored at -20°C until day of injection and diluted to 1% in sterile saline. Retrogreen beads were purchased from LumaFluor (Durham, NC [Cat. Retrobeads IX]) and stored at 4°C. Full details of each tracer can be found Table 2.

**Table 2.**

| Tracer Name | Chemical Class | Key Properties | Manufacture product # | Observations | Rating (1-4) |
| --- | --- | --- | --- | --- | --- |
| **Fluoro-Gold (FG)** | Fluorinated stilbene derivative | High sensitivity, stable, long-term labeling | Fluorochrome Fluoro-Gold 20mg | Good but low contrast | ★ ★ ★ |
| **Fast Blue (FB)** | Stilbene derivative | Excellent uptake, strong fluorescence, long retention | Polysciences 17740 | Robust fluorescence | ★ ★ ★ ★ |
| **True Blue (TB)** | Stilbene derivative | Similar to Fast Blue with lower background | MedChem Express HY-D1161 | Robust fluorescence | ★ ★ ★ ★ |
| **Cholera Toxin B (CTB)-647nm** | Protein (non-toxic subunit B) | Bidirectional; uptake via GM1 gangliosides | ThermoFisher Scientific C34778 | Non-specific retrograde labeling | ★ ★ |
| **Retrogreen Latex Microspheres** | Xanthene derivative | Robust fluorescence, no leakage at injection site | LumaFluor Green Retrobeads™ IX | Dull signal detected | ★ |

### Retrograde tracer Intra-articular injections

Retrograde tracers were injected intra-articularly in 2-month-old wild-type mice. The sample size for each retrograde tracer was three (N = 3). Briefly, mice were placed under anesthesia using 2-3% isoflurane, and the skin overlying the knee was depilated and disinfected with three applications of diluted betadine and 70% alcohol, prior to injections. In a single intra-articular injection, four microliters of a retrograde tracer were delivered to a single mouse using a 30G needle attached to a 100 μl Hamilton glass syringe. Retrograde labeling studies were terminated 5 days after injections.

### Tissue collection and processing

Mice were euthanized using CO_2_ and cardiac perfused with 15 mL of cold 1 X PBS followed by 15 mL of 4% PFA (Thermofisher; Waltham, MA [Cat. J19943.K2]. Following perfusion with 4% PFA in 1 X PBS, the spine was resected by removing hindlimbs, cutting away internal organs and making perpendicular cuts at cervical and sacral vertebrae. Mouse knee joints were harvested, fixed in paraformaldehyde, and decalcified for coronal embedding in optimal cutting temperature (OCT) media as previously described^42^. Discrete bilateral sympathetic chain ganglia were post-fixed in 4% PFA in PBS for 1-3 hours at 4°C, washed in 1 X PBS and placed in 30% sucrose (Thermofisher; Waltham, MA [Cat. A15583.36]) overnight, at 4°C. Ipsi- and contra-lateral lumbar (L3-L5) DRGs were identified and processed separately, similar to sympathetic chain ganglia tissues. Fixed neuronal tissues were embedded in Tissue Tek-O.C.T compound (VWR North American [Cat: 4583]), frozen at -20°C and sectioned using a cryostat (Leica Nussloch, Germany). Chamber temperature of cryostat was set to -20°C and DRG tissues were sectioned at 20 microns; sympathetic ganglia chains were sectioned at 20 microns. All tissue was mounted on Superfrost microscope slides (VWR North American [Cat: 48311-703]) and allowed to adhere to the slide at room temperature for 10 minutes, prior to storing at -20°C.

### Immunofluorescence

Knee sections (25 µm) were collected and stained with an anti-βIII tubulin antibody (Abcam, [ab18207]) at a 1:200 dilution, following the same protocol^42^. A donkey anti-rabbit Alexa Fluor 647 secondary antibody was used at a 1:700 dilution. Twenty micron sections of DRG and SG were probed for neuronal markers similar to previously published methods^43^. Briefly, primary antibodies raised against NeuN (1:500), (Abcam; Cambridge, UK; [Cat:177487]) Tryosine hydroxylase (1:200), (Millipore; Burlington, MA; [cat: ab152]) were diluted in a buffer containing 1 X PBS, 3% Bovine Serum Albumin [Cat: 9048-46-8], 5% normal donkey serum (Jackson ImmunoResearch; Chester County, PA [017-000-121] and 0.3% triton-X 100 (Millipore; Burlington, MA [Cat: 9036-19-5]); and incubated overnight at 4°C. Negative control samples were incubated in buffer without primary antibody. Probed tissues were washed in 1 X PBS and 0.1% Triton X 100, 3 times, 5 minutes each wash. Alexa-Fluor 488 (Thermofisher; Waltham, MA; [Cat: A-21206]) or 594 ([Cat: A-21207]) secondary antibodies raised against rabbit IgG were diluted 1:250, in buffer and incubated with samples for 2 hours at temperature. Samples were washed and stained with either DAPI (1:4,000) (Thermofisher; Waltman, PA [Cat: 62248] or ToPro3-647 (1:1,000) (Thermofisher; Waltman, PA [Cat: T3605]) nucleic acid dyes. Samples were washed in 1 X PBS 3 times and mounted with ProLong Gold mounting medium (Thermofisher; Waltman. PA [Cat: P36934]) and a coverslip and allowed to cure overnight at room temperature. Specificity of reporter (TdTom) expression patterns was confirmed by subjecting DRG and SG sections, obtained from reporter mice (Ai9 or Ai65F) not carrying recombinase genes, treated to the above-described immunofluorescent procedure in the absence of primary antibody (negative control), as shown in Figures 1-3 and Supplemental Figure. 4.

**Figure 1.**
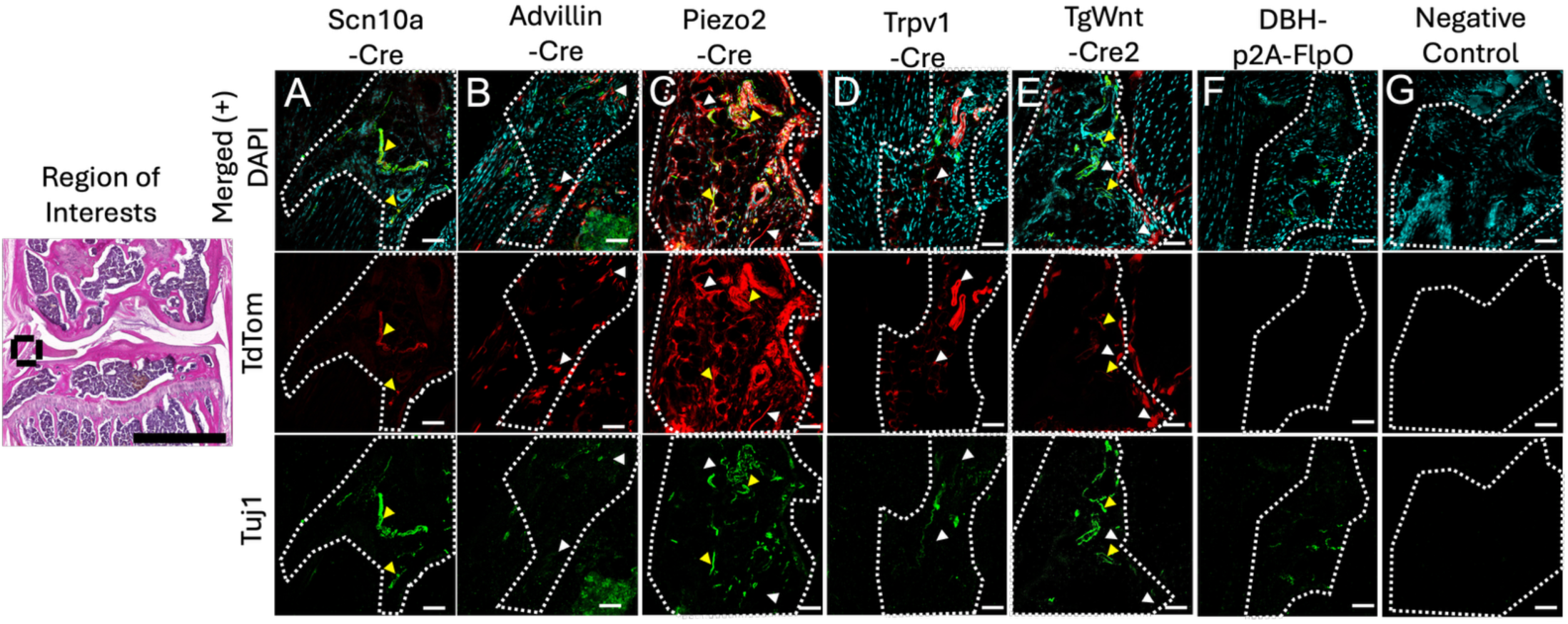
Representative images taken of the synovial membrane (area within dotted lines) from middle mid-coronal knee sections of genetic reporter mice. Merged channels in the first row represent pattern of TdTom and Tuj1 colocalization. White arrows were overlayed on to images to indicate lack of Tuj1 colocalization with TdTom expression in merged, TdTom and Tju1 channels. Colocalization of Tuj1 and TdTom expression was labeled with yellow arrow heads. Every column represents a different genetic reporter screened in this study. Similar Knee sections not treated with primary antibody were taken from Ai9 or Ai65F (+) mice (no driver) as “double” negative for Cre recombinase and primary Tuj1 antibody (G).

**Figure 2.**
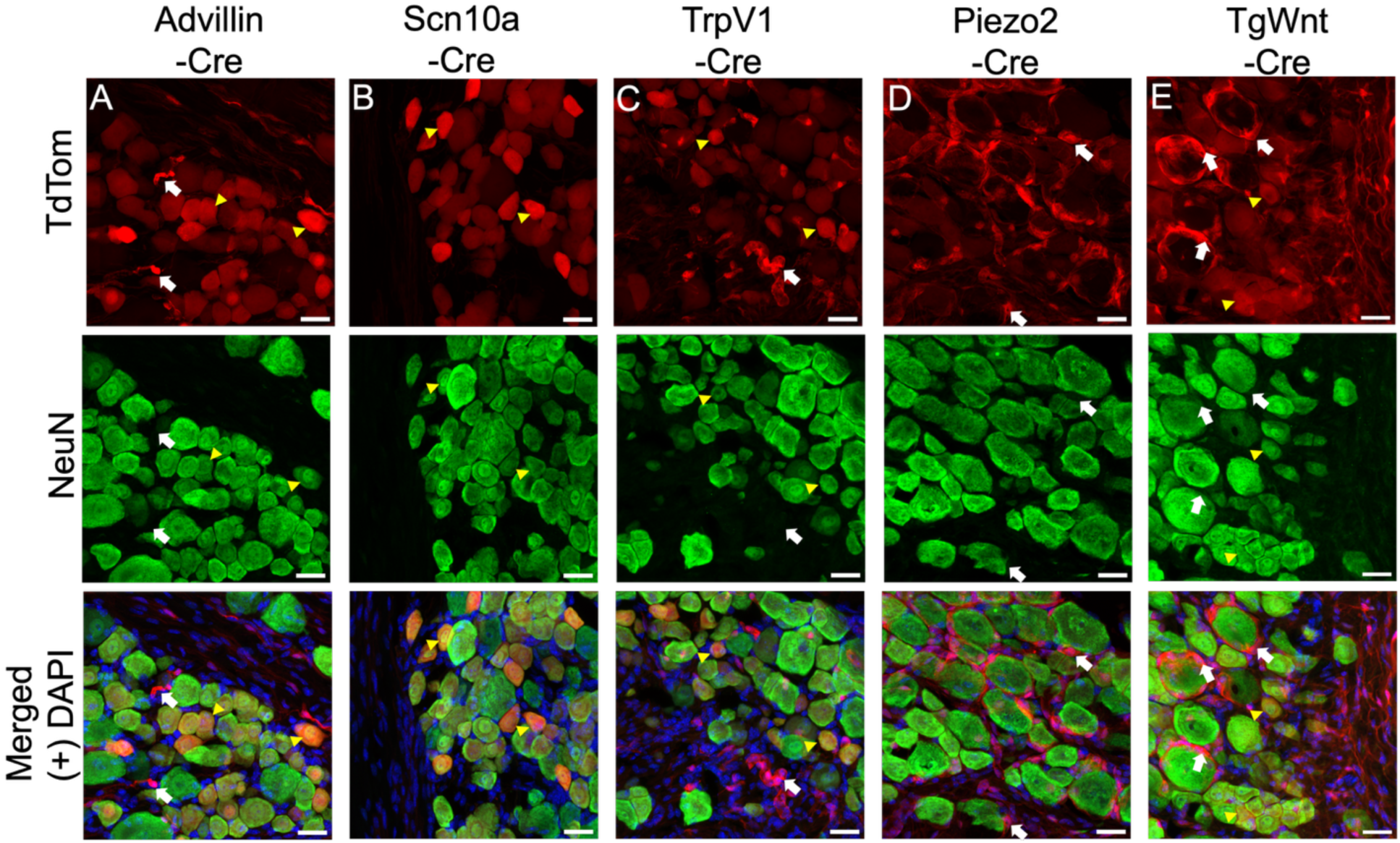
Representative images of DRGs taken from genetic reporter mice. Neuronal expression of TdTom signal was confirmed by staining with neuronal nuclei marker NeuN. Genetic reporter expression in Advillin-Cre (A), TrpV1-Cre (C), PIEZO2-Cre (D) and TgWnt-Cre (E) was observed in neurons and no-neuronal cells. White arrows point to areas of TdTom expression absent of NeuN colocalization. Yellow arrow heads point to cells with reporter and NeuN colocalization. The Scn10a-Cre (B) reporter mouse model exclusively expressed TdTom signal in NeuN expressing cells. Images are Z-projections spanning 20 microns. Scale bar = 25 microns.

**Figure 3.**
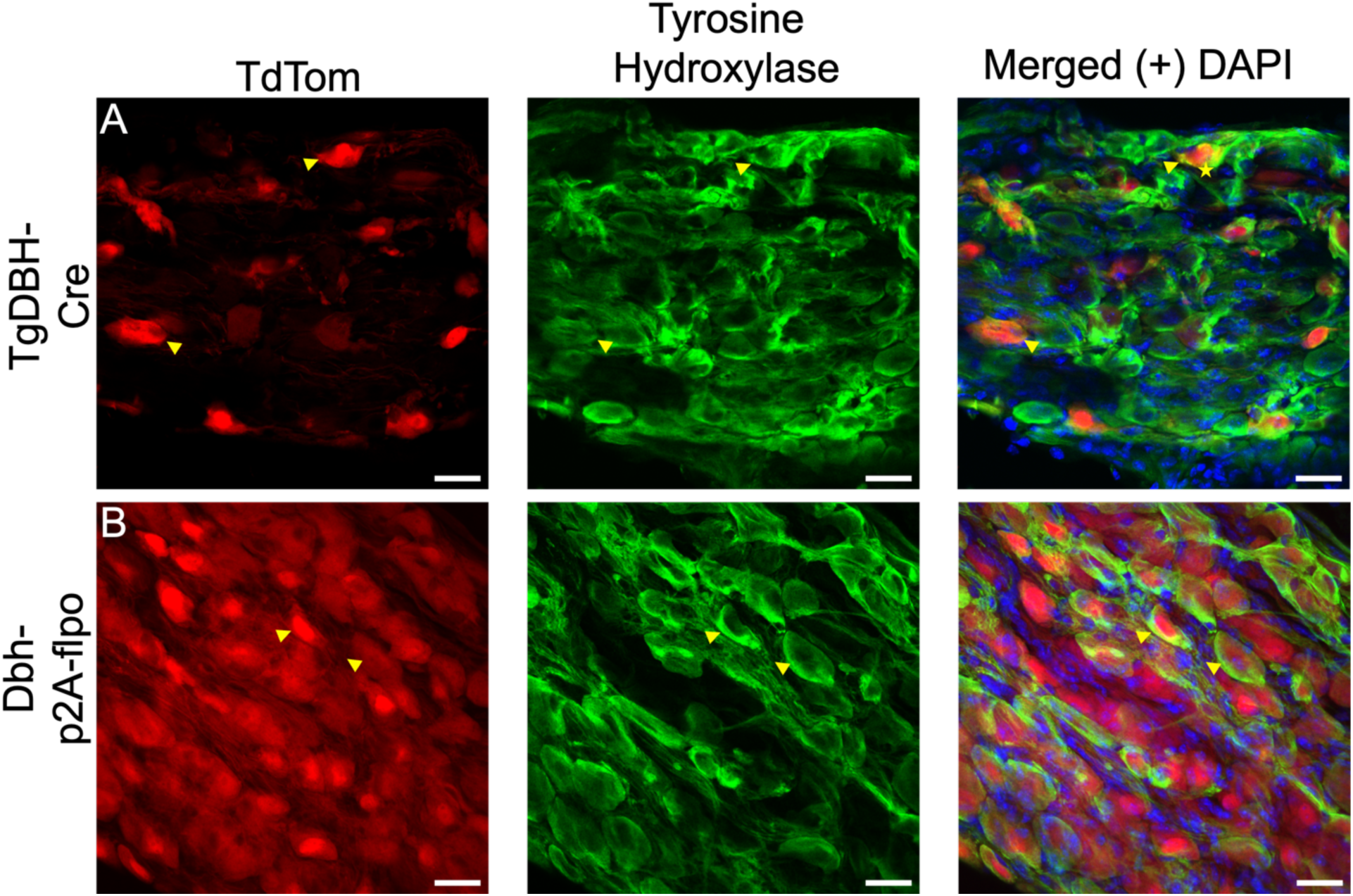
Representative images of sympathetic chain ganglia from TgDBH-Cre (A) and DBH-FlpO reporter mice (B). Specificity of sympathetic TdTom signal (red psuedocolor) was confirmed by staining with Tyrosine Hydroxylase antibody (green psuedocolor). Images are Z-projections spanning 20 microns Scale bar = 20 microns.

**Figure 4.**
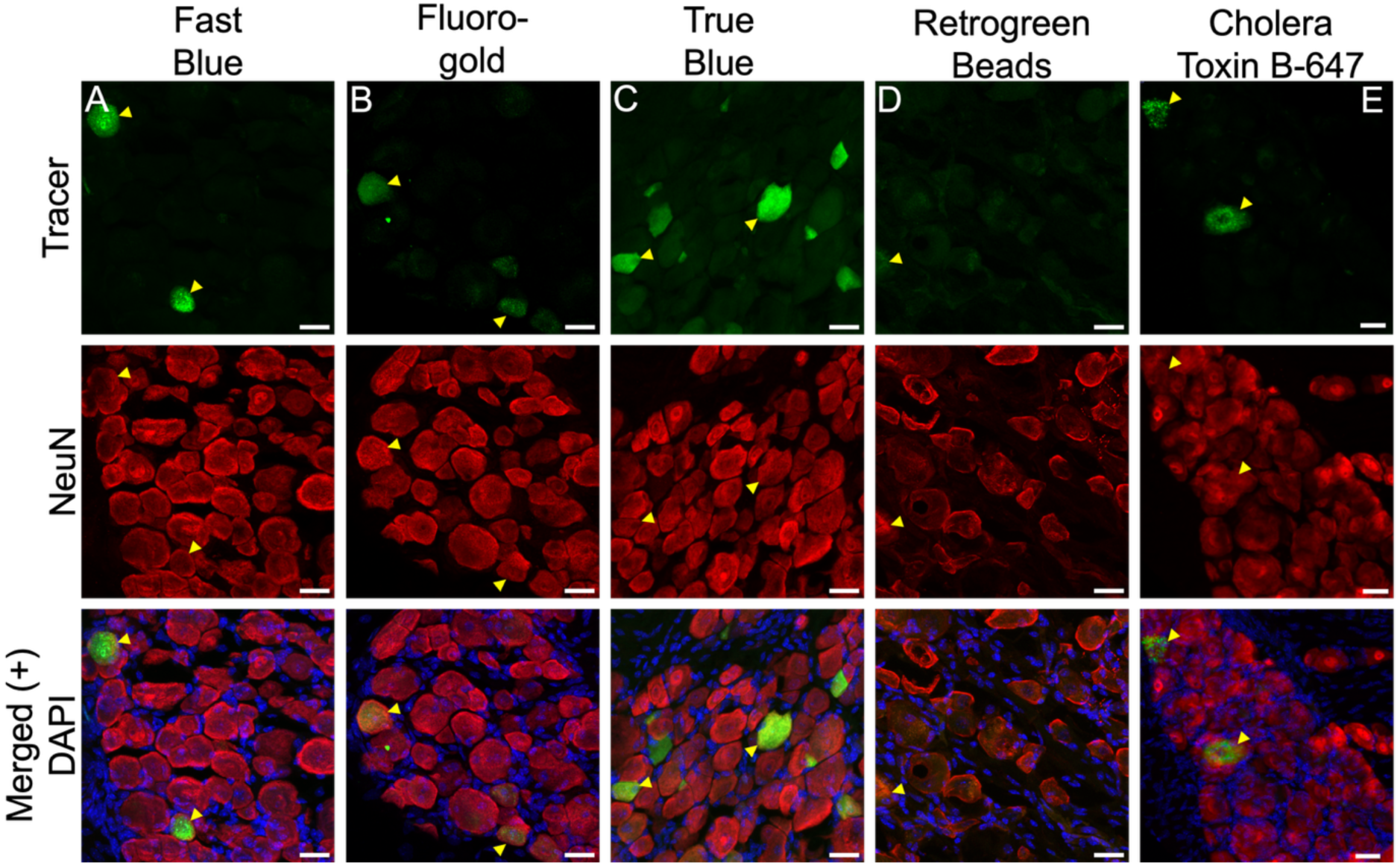
Representative images of DRGs collected from wild type mice injected intra-articularly with conventional retrograde tracers. DRG sections were stained labeled with neuronal nuclei marker NeuN (pseudocolored in red). Conventional retrograde tracer Fast Blue (A), Fluorogold (B), True Blue (C), and Cholera Toxin B-647 (E) are represented in green (psuedo) color. The Retrogreen Beads tracer (D), was not observable using similar adjustments to brightness settings. Yellow arrow heads point to tracer and NeuN colocalization. Images are Z-projections spaninng 20 microns. Scale bar = 25 microns.

### Microscopy and Image analysis

For Knee samples, confocal imaging was performed using a Zeiss LSM 880 Airyscan FAST microscope with a 20x objective. Z-stacks were acquired at 1 µm intervals to a total depth of 16 microns. Ganglia samples were captured on the same microscope without Airyscan FAST capture enabled. All representative images were processed using Fiji software by creating z-projections with max intensities from original Zeiss files. Retrograde tracers were captured and pseudocolored green for a consistent appearance in Figure 4. Confocal microscope and Fiji imaging software settings were maintained between bioreplicates for each reporter mouse line and retrograde tracer. In Fiji imaging software, moderate adjustments to brightness and contrast were performed to increase brightness over entire sample in representave images.

## Results

### Neural characterization of genetic recombinase mouse lines in healthy knee joint tissues

We surveyed reporter gene expression in intact knee joints, focusing on the mid-region of the knee where both cruciate ligaments can be observed in the same plane(**Fig. 1**). All sensory neuronal reporter lines showed crisp, sharply outlined TdTom-positive cells that resemble neurons in the lateral synovium, while reporter expression levels were below detectable limits for the sympathetic driver (**Fig.1).** We next co-stained these sections for the pan-neuronal marker beta-3 tubulin (Tuj1) to confirm the identity of TdTom-expressing cells. We found that most Cre lines induced extensive recombinase in non-neuronal cell types in this tissue, as determined by the absence of Tuj1 staining in TdTom-positive cells (**Fig. 1**, white arrows). The Scn10a-Cre nociceptor reporter model (NaV1.8) showed crisp signal in synovial sections that exclusively colocalizes with Tuj1 nerves (**Fig. 1A**, yellow arrows). In the pan-neuronal Advillin-cre reporter, the strongest TdTom signal did not colocalize with Tuj1 staining, and there was minimal overlap between both markers in healthy synovium (**Fig. 1B**). Widespread fluorescent signal from the somatosensory model Piezo2-cre mice was observed in Tuj1 positive nerves, though this line was also strongly expressed in non-neuronal synovial tissues (**Fig. 1C**). While the somatosensory reporter line (TrpV1) exhibited localized expression in DRG neurons, we observed very little co-localization in knee sections (**Fig. 1D**). For Wnt1-Cre, TdTom signal was widespread and strong in cells that did not co-stain with Tuj1 (**Fig. 1E**). Lastly, the sympathetic reporter line (DBH-p2A-FlpO) did not show detectable reporter expression in the healthy synovium (**Fig. 1F**). To survey innervation throughout the knee joint, we assessed reporter gene expression in three additional regions (cruciate ligaments, medial synovium, subchondral bone). Compared to the lateral synovium, the medial synovium, cruciate ligaments and subchondral bone revealed very limited sensory innervation, and fluorescent signal (**Suppl. Fig. 2, 3**). In sum, sensory genetic drivers nominally expressed fluorescent reporter in neuronal tissues that permeate the healthy, adult knee joint, though expression was limited and more likely to highlight non-neuronal tissues. In contrast, the DBH-p2A-FlpO reporter did not express observable fluorescent signal in any knee joint tissues.

### Neural characterization of genetic reporter expression in sensory and sympathetic ganglia

We observed reporter gene (TdTom) expression in DRGs obtained from all sensory neuron drivers, and in SGs from sympathetic drivers (**Fig. 2, 3).** Of note, we did not detect any TdTom expression in DRGs of sympathetic drivers or in SGs obtained from sensory neuronal drivers (Data not shown). To confirm the specificity of selected driver lines, we co-stained DRG sections with an established neuronal marker-NeuN, (**Fig. 2)**. Importantly, we detected extensive non-neuronal TdTom expression in several sensory driver lines (**Fig. 2**, white arrows). Advillin-Cre expression was widespread in neurons (yellow arrows) and in smaller non-neuronal cells (Fig 2 A). Scn10a-Cre (NaV1.8) mice expressed robust TdTom signal that co-localized with NeuN in DRG sections (**Fig. 1B**). The somatosensory driver Trpv1 expressed strong TdTom signal in neuronal and non-neuronal cells (**Fig. 2**C). The Piezo2 reporter expressed the strongest signal in non-neuronal cells surrounding NeuN-positive neurons (**Fig. 2**D). The TgWnt1-Cre reporter induced widespread signal in neuronal and non-neuronal fluorescent in DRG sections (**Fig. 2**E). Hence, only NaV1.8 reporter mice displayed specific reporter gene expression in sensory neurons in the DRG only, while all other sensory reporter lines tested showed some additional non-neuronal reporter gene expression. Genetic driver lines in this study were summarized and rated for their specificity to label knee innervating neurons (**Table 1**). Transgenic DBH-Cre recombinase was chosen to compare transgenic and knock-in sympathetic drivers^44^. Upon crossing selected driver lines to a Cre- or Flp-responsive fluorescent reporter line (Ai9 or Ai65F), SG were collected and stained for tyrosine hydroxylase (TH) to mark sympathetic neurons (**Fig. 3**). In contrast to available tools to study sensory innervation, the sympathetic driver lines tested here showed exclusive reporter gene expression in sympathetic neurons, as visualized by colocalization (yellow arrows) with the sympathetic neuron marker, tyrosine hydroxylase (TH) (**Fig. 3**). When comparing expression patterns between both selected sympathetic driver lines, the knock-in DBH-p2A FlpO (A) reporter mouse showed more robust and widespread sympathetic neuronal signal compared to the TgDBH-Cre reporter (B).

### Evaluation of retrograde tracer dyes

Retrograde tracers (**Fig. 4)**. Fast Blue (**Fig. 4A**) and True Blue (**Fig. 4C**) showed robust and unambiguous fluorescent signal, consistently filling neuronal soma bodies throughout the DRG. The fluorogold retrograde tracer (**Fig. 4B**) filled neuronal soma, but displayed lower signal intensity and appeared in fewer DRG sections. In contrast, labeling joint-innervating neurons with Retrogreen Beads fluorescent dye (**Fig. 4D**) resulted in weak neuronal labeling in the DRG that was difficult to detect above background fluorescence. Lastly, while the fluorescent signal from Cholera Toxin B-647 (**Fig. 4E**) filled neuronal soma in DRGs collected from the same side as the intraarticular injection, this tracer could also be detected in contralateral DRGs from uninjected knees, rendering this tracer less suitable for neuronal circuit mapping studies in the knee(**Suppl Fig. 2, Fig 2**). The performance of these retrograde tracers were evaluated and rated for their performance and specificity to label knee innervating neurons (**Table 2**).

## Discussion

Musculoskeletal pain, particularly related to joint diseases such as osteoarthritis and rheumatoid arthritis, forms a significant global health problem that is only expected to grow^6,45^. As opiates are increasingly used in treating OA pain, this creates an additional pathway the growing opiod overdose epidemic. To mitigate the use of opiates and to clear a path towards improved treatments for joint disease progression and pain management, the NIH established the RE-JOIN consortium to build comprehensive anatomical and molecular maps of the knee and temporal mandibular joints.

While numerous studies in the last century have described joint innervation, they typically focused on individual tissues or neuronal subtypes using limited tools including immunohistochemistry. Hence, whole-joint innervation maps that reveal the intricate network of distinct subtypes of sensory or sympathetic neurons and the musculoskeletal tissues they permeate are sorely lacking. Given recent studies revealing excessive nociceptive sprouting in human and animal models of OA^46–48^, a comprehensive neural map of the knee joint beyond nociceptive fiber types is needed to better understand the neurobiology of joint pain and reveal possible neuro-specific therapeutic targets. However, histological analysis of joints is notoriously challenging, as this organ is composed of several mineralized and non-mineralized tissues, requiring specific processing steps that increase tissue autofluorescence and affect epitope integrity^39^. These challenges, combined with low specificity of antibodies for membrane proteins, and thinly branched neuronal morphology so far have limited researchers to study joint-innervating populations by fiber type. Further, these histological methods often lack subtype resolution and preclude the ability to molecularly profile and functionally interrogate targeted populations. To overcome these challenges, researchers have turned to genetic recombinase mouse lines combined with fluorescent tracer dyes to map population and projection specific neural networks. However, numerous studies have shown that care must be taken when using Cre and Flp recombinase lines, as they may yield expression in fewer or more cell types than those they were designed to target and importantly represent a cumulative fate map of expression that occurred at any time in the animal’s life, irrespective of how transient^49^. In addition, scientists have studied tissue innervation using chemical retrograde tracers, but effective labeling in the joint can be hampered by the heterogeneous nature of synovial joint tissues, sparse innervation, and tracer leakage into muscle and fatpad^50^. Here, we addressed some of these challenges by developing a validated toolkit of genetic and molecular resources to map knee joint innervation in the mouse. To do so, we performed in-depth expression analysis in the knees, DRGs, and SGs of sensory and sympathetic driver lines to identify those that reliably label the expected neuronal subtype. In addition, we tested several classes of conventional retrograde tracers to evaluate their efficacy in labeling knee-innervating neurons. To our knowledge, this type of comprehensive comparative analysis to validate commonly used neural labeling reagents has not been conducted to date. Our results showcase the importance of screening genetic reporters for neural specificity and of identifying specific retrograde tracers for robust circuit mapping in the knee.

The majority of sensory neuronal driver lines used in this study displayed extensive Cre-activity in both neuronal and non-neuronal tissues, limiting their utility. Although genetic neural reporters were selected based on their reported primary role in peripheral neurons, they exhibited substantial reporter signal in non-neuronal tissues throughout the knee joint and DRG. For some lines (Piezo2-Cre, TrpV1-Cre, Wnt1-Cre2), some non-neuronal expression was expected in the knee based on previous reports^29,32,51–53^. Furthermore, transgenic recombinase lines often show expression patterns that differ from their target gene due to random transgene insertion in to the genome and other design-related consequences. It was surprising to observe non-neuronal Cre activity in the knee of the Advillin-cre mouse line since it was originally reported to be primarily expressed in peripheral neurons^28^. Advillin is an actin binding protein involved in sensory axonal growth during development and nerve injury^28^. We report that the Advillin-Cre recombinase driver line expressed robust signal in non-neuronal tissues in knee and DRG sections (Figures 1-B, 2-A).

The TrpV1 receptor is important for nociception and is activated by noxious heat stimuli^54,55^. Mapping TrpV1-expressing neural subpopulations innervating the knee would allow us to visualize changes in thermal sensing neural networks associated with OA^56,57^. In the knee joint and DRG, we detected extensive genetic reporter expression in non-neuronal tissues that are consistent with expression patterns observed by Cavanaugh et al., 2011^29^,. The Piezo2 receptor plays a role in non-nociceptive and nociceptive mechanical sensation^58^. Early studies demonstrated that Piezo2 is activated by light touch, but more recent investigations found decreased mechanical allodynia from acute inflammation in the absence of Piezo2 receptor, suggesting Piezo2 is involved in relaying nociceptive signals ^30,59,60^. Given that OA patients who report mechanical sensitivity are more susceptible to developing chronic pain, the ability to map joint disease-induced changes to Piezo2^+^ knee-innervation would enable us to study how these changes correlate with pain behaviors and improve our understanding of pain chronification in the context of OA^61^. However, we observed widespread and robust reporter gene expression in non-neuronal tissues across knee and DRG sections of Piezo2-Cre mice. Recent studies have started to investigate the function of Piezo2 outside of the peripheral nervous system, and have reported low-to-medium Piezo2 mRNA expression in bone (osteoblasts), cartilage, and synovial blood vessels^62,63^. Hence, studying this neuronal subtype in the knee and DRG will require co-labeling for a neuron-specific marker, such as Tuj1 or NeuN. In addition, further studies will be needed to reveal whether the non-neuronal Cre-activity we detected in the DRGs of Piezo2-Cre mice is an artefact specific to this Cre line, or if Piezo2 plays a thus far unknown role in this cell type and at what stage of development or aging^53,62,63^. The sodium channel, Scn10a/Nav1.8 is a voltage gated channel found primarily in small-diameter nociceptor neurons and is important for mediating mechanical sensation and hyperalgesia^64^. Furthermore, the Nav1.8 channels are important targets for chronic pain but do not mediate initial pain signals. Instead they are expressed in response to persistent pain signaling from somatosensory channels^57,65^ In particular, previous OA studies have shown changes in nociceptive neural networks in OA knees using antibody-based detection methods as well as this Scn10a-Cre reporter model^46,47^. Our results confirm that this mouse line is a reliable tool for mapping knee-innervating somatosensory neurons without the need for additional labeling, as we detected robust reporter gene expression exclusively in sensory neurons in the knee and DRG. Sympathetic neurons are integral for joint health and highly represented in the knee^37,38,66^. Early retrograde tracing studies demonstrated that a significant proportion of knee innervating neurons originate from the lumbar sympathetic chain ganglia^37,38^. Further investigation into their role in the knee showed that noradrenergic neurons are crucial for knee joint homeostasis through regulation of bone and cartilage growth^66,67, 68^. Joint-disease-induced changes in sympathetic distribution have been observed in synovial tissues of patients with rheumatoid arthritis, and in preclinical models of temporomandibular joint disorders, suggesting dynamic sympathetic joint innervation in the context of joint pain^69,70^. Similarly, studies in murine models of traumatic nerve and tissue injury show that sympathectomy or adrenergic blockade can reduce chronic pain^71,72^. To be able to visualize this important neural network in the knee, we screened two different sympathetic DBH reporter models. In both genetic DBH Cre and FLPo based reporter models, we observed exclusive noradrenergic neuron colocalization with TH staining. Though, the expression pattern in the transgenic TgDBH-Cre reporter was much less robust than the knock-in DBH-p2A-FlpO reporter model with fewer cells expressing TdTom. The DBH-p2A-FlpO model was first created by Sun et al., 2017 and exhibited robust colocalization of noradrenergic neurons in the brainstem while preserving DBH expression^24^. Consistent with their findings, our study was able to demonstrate robust and exclusive reporter expression in noradrenergic neurons for this driver. Although, *DBH* driven expression of the reporter was not detected in knee sections, robust and specific labeling of noradrendergic cell bodies in chain ganglia was observed in both sympathetic reporter models, indicating further optimization is required to detect sympathetic nerve endings in target tissue. Previous antibody-based studies reported tyrosine hydroxylase-positive neurons innervating healthy and arthritic knees^69,70^. However, the TH antibody used for these studies labels both sympathetic neurons and a subset of sensory neurons. Hence, the nature of the TH-positive neural network permeating the knee joint remains unclear. To date, retrograde labeling of sympathetic neurons is the strongest evidence of sympathetic knee innervation and therefore it may be necessary to enhance detection of reporter signal using tyramide signal amplification in the DBH-p2A-FlpO reporter model.

There are some limitations to this study to consider, including the number of tissues analyzed in this study that are often studied in the context of joint diseases (synovium, subchondral bone). Additionally, this study characterized TdTom expression from healthy joint tissues which could change in the context of joint disease, like post-traumatic osteoarthritis (PTOA) mouse models including Anterior Cruciate Ligament Tear (ACLT) and Destablization of Medial Meniscus (DMM) models. This study was able to characterize two mouse lines with neuron-specific reporter expression which can be used in tissue clearing and light-sheet microscopy. This application could overcome some of the challenges encountered when using traditional histological methods in 2D, and provide a higher resolution, 3D map of subset neural networks and their inter-connections with other neurons and cell types throughout the entire knee.

The diversity of retrograde tracer dyes employed thus far for tracking knee innervating neurons are wide-ranging and yet there is no consensus on which tracer class is optimal^35^. In this study, we evaluated several popular classes of retrograde tracer dyes and found that stilbene-derived tracer dyes provide consistent and robust neuronal signal without leakage. Fast Blue and True Blue performed the best in tracing knee innervating neurons; showing robust somatic neuron labeling. Fast Blue is widely used for mapping the origin of peripheral neurons due to its easy preparation and robust labeling of sensory neurons^35^. Delivering consistent labeling patterns, Fast Blue is commonly used as a positive control to optimize other retrograde tracers^,35,50,73^. More recently, Fast Blue was used to study *in vivo* calcium responses of knee innervating neurons in the context of OA^74^. However, in our experience, commercial availability of Fast Blue due to supply chain limitations has been an issue. While it is unclear if shortages will persist in the future, it may be worth considering alternative approaches, including viral tracers for long-term studies. Although previous studies report similar characteristics, True Blue is an underutilized retrograde tracer and has not been reported for mapping the origin of knee innervation neurons thus far. Our study now shows that True Blue-labeled neurons emitted robust fluorescent signal without leakage into neighboring cells. Hence, True Blue serves as a reliable alternative to Fast Blue for labeling knee joint-innervating neurons. The fluorinated stilbene derivative, Fluorogold labeled knee innervating neurons but resulted in a weaker signal compared to Fast Blue and True Blue. The low contrast from Fluorogold is consistent with previous studies but remains viable for tracing peripheral neurons given its resistance to photobleaching and antibody enhancement^35^. Cholera Toxin B-647 clearly labeled neurons as expected but was quickly transported to the contra-lateral DRG side in this study. This result is consistent with a previous report where biotin-CTB (similar uptake mechanism to CTB-647) was also observed on the contra-lateral side of sciatic nerve injections^75^. Cholera Toxin B-Alexa Fluor is a popular choice for retrograde tracing given its precise mechanism of transport, fast transport and robust neuron labelling. Therefore, the utility of CTB-Alexa Fluor will depend on study design and need for internal controls. Lastly, Latex Microspheres have become another popular choice for neural circuit tracing due to benefits including resistance to fading, photobleaching, and diffusion at the site of injection field^35^. Expecting results similar to Silva Serra et al., 2016, where microbeads were used to compare joint and cutaneous innervating neuron populations, we did not detect knee innervating neurons with several intra-articular (Fig. 4D) and intra-epidermal (data not shown) injections with Green Retrobeads^50^. While it is possible that Green Retrobeads do not transport similarly to the Red Retrobeads used in this paper, we did not test this hypothesis in the current study.

Here, we established an important toolkit for labeling joint-innervating neurons by comparing popular conventional retrograde tracers and evaluating their signal in primary afferent neurons. We did not test another important tool, viral retrograde tracers but efforts to employ this technology to map neural subpopulations need to be addressed in future comparative studies.

In conclusion, screening popular mouse reporter strains and retrograde tracers revealed that very few genetic drivers exclusively express fluorescent reporter in neurons and that stilbene derivative tracer dyes were an optimal tracer dye for tracing knee innervating neurons. Our recommendation for comprehensive mapping of synovial joints is to employ the Scn10a-cre driver with Ai9 reporter strain and DBH-p2A-FlpO driver with Ai65F reporter strain in combination with the retrograde tracer dye, True Blue. This study is the first to report a large-scale characterization of multiple genetic reporters and retrograde tracers for mapping knee innervating neural circuits. The results of which provide an invaluable resource for researchers investigating peripheral nerves in musculoskeletal related diseases like RA and OA.

There are several notable applications for this toolkit in building multifaceted neuronal maps of knee joint innervation. First, we utilized both Cre and FLP recombinase systems. This enables the future application of intersectional genetics, whereby two separate recombinases are used in the same mouse, i.e., both Cre and FLP drivers, with a specially designed reporter allele that turns on only in neuronal populations that express not just one genes specific recombinases but both recombinases in the same cell^25,76,77^. This offers even higher resolution as markers characterized in this paper are known to mark neurons that show further genetic heterogeneity within their defined sensory and sympathetic populations. Genetic expression of a fluorescent protein not only facilitates anatomical mapping and visualization, but it also creates access to these neurons for unambiguous electrophysiological characterizations and isolations of neurons that can then be used for single-cell/nuclear RNA sequencing and proteomics^78^. Lastly, a large number of single and double recombinase responsive alleles exist that express a number of optogentic and chemogentic effector molecules that allow for *in vivo* functional characterizations in awake and freely behaving animals^79^. Lastly, by layering retrograde tracers over the genetically delineated neuronal populations, an additional spatially defined layer of resolution is achieved, whereby genetically defined neurons are further subdivided by their projection targets. Though this resolution can be leveraged for anatomical and molecular studies, it does not lend itself to functional manipulations. Such spatial or projection-target resolution for functional manipulations may be achieved by expressing a heterologous viral receptor in a recombinase-dependent fashion and using viral injections to express an optogenetic or chemogenetic effector molecule for functional interrogations of target neurons in disease progression and pain development^80^. Altogether, this work, creates a rich foundation for further characterizations of the knee innervating neurons in healthy and disease models that will yield important clues and clear a path to therapeutic interventions to improve disease progression and improve pain outcomes, contributing to the RE-JOIN goal of improving patient outcomes and reducing the use of opiates that may contribute to the current opiate overdose epidemic.

## Author contributions

Ibdanelo Cortez, Nele A. Haelterman, Brendan H. Lee, and Russell S. Ray conceptualized and oversaw the study. Ibdanelo Cortez (Sympathetic ganglia), Vanessal Belizaire (Dorsal root ganglia), and Carolina Leynes (Knee joints) executed tissue dissections, and stained tissues. Ibdanelo Cortez (ganglia tissues) and Carolina Leynes (Knee joints) captured and processed images. Brendan H. Lee and Russell S. Ray obtained funding and provided lab resources for the study. Ibdanelo Cortez, and Nele A. Haelterman wrote the original draft. All authors revised the paper and consented to its contents.

## Acknowledgements

The authors are grateful to the members of RE-JOIN consortium at Baylor College of Medicine for contributing their expertise and valuable discussions related to this work, and for their insightful comments and edits. This work was supported by the National Institutes of Health through the Helping to End Addiction Long-term Initiative and the National Institute of Arthritis and Musculoskeletal and Skin diseases (UCAR082200). This paper’s content is solely the responsibility of the authors and does not necessarily represent the official views of the National Institutes of Health.

The RE-JOIN consortium consists of: Armen Akopian, Kyle Allen, Alejandro Almarza, Benjamin Arenkiel, Basak Ayaz, Yangjin Bae, Bruna Balbino de Paula, Anita Bandrowski, Mario Danilo Boada, Jacqueline Boccanfuso, Jyl Boline, Dawen Cai, Carpio, Dellina Lane, Robert Caudle, Racel Cela, Yong Chen, Rui Chen, Brian Constantinescu, Cortez, Ibdanelo, Yenisel Cruz-Almeida, M. Franklin Dolwick, Chris Donnelly, Zelong Dou, Joshua Emrick, Malin Ernberg, Danielle Freburg-Hoffmeister, Spencer Fullam, Janak Gaire, Akash Gandhi, Benjamin Goolsby, Stacey Greene, Nele Haelterman, Michael Iadarola, Shingo Ishihara, Azeez Ishola, Sudhish Jayachandran, Zixue Jin, Frank Ko, Priya Kulkarni, Zhao Lai, Brendan Lee, Yona Levites, Carolina Leynes, Jun Li, Martin Lotz, Lindsey Macpherson, Tristan Maerz, Camilla Majano, Anne-Marie Malfait, Maryann Martone, Bella Mehta, Richard Miller, Rachel Miller, Michael Newton, Alia Obeidat, Merissa Olmer, Dana Orange, Miguel Otero, Kevin Otto, Folly Patterson, Marlena Pela, Sienna Perry, Theodore Price, Hernan Prieto, Russell Ray, Dongjun Ren, Margarete Ribeiro Dasilva, Alexus Roberts, Elizabeth Ronan, Oscar Ruiz, Shad Smith, Mairobys Soccorro Gonzalez, Kaitlin Southern, Joshua Stover, Michael Strinden, Hannah Swahn, Evelyne Tantry, Sue Tappan, Luis Tovias Sanchez, Airam Vivanco-Estela, Joost Wagenaar, Lai Wang, Kim Worley, Joshua Wythe, Jiansen Yan, and Julia Younis.

## Supplemental Figures

**Supplemental Figure 1:**
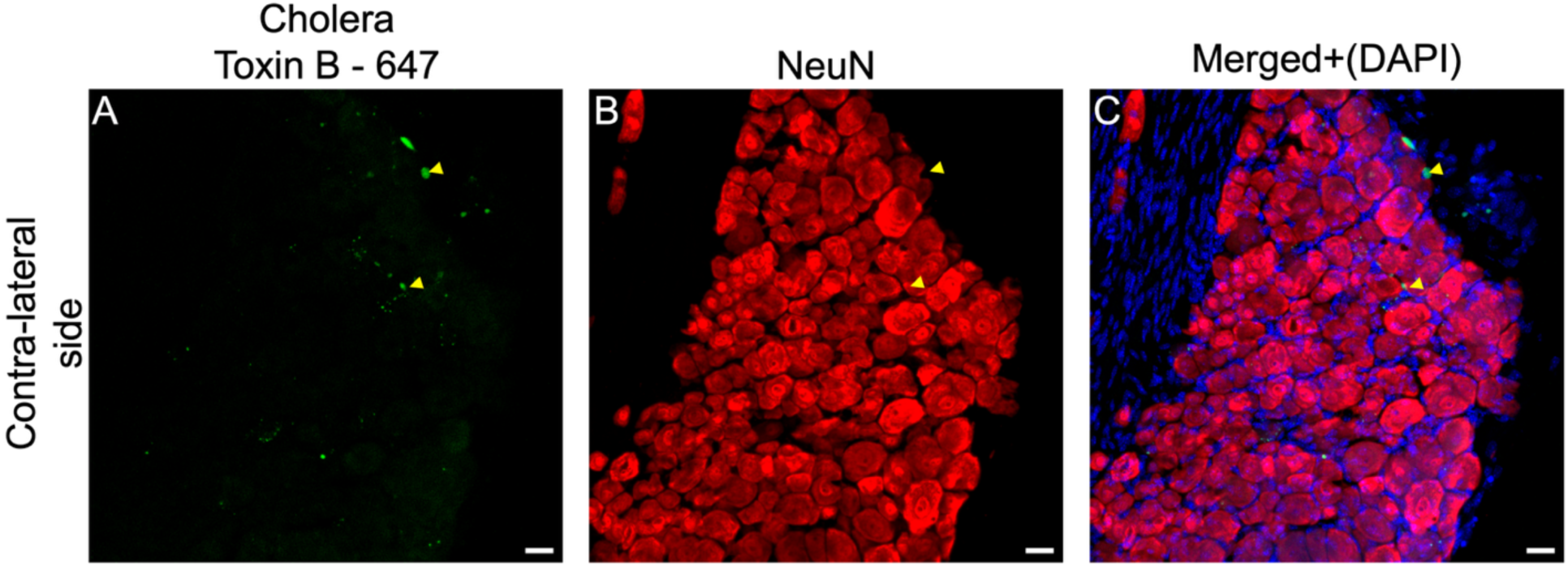
Representative Images captured of contra-lateral DRG taken from mouse injected with CTB-Alexa Fluor 647. Panel A shows positive signal from CTB-647 tracer in green (psuedocolored). DRG section was stained with NeuN (psuedocolored in red), panel B. Merged channels with nucleic acid stain, DAPI panel C. Yellow arrow point to cells with tracer and NeuN. Images are Z-projections spanning 20 microns. Scale bar = 25 microns.

**Supplemental Figure 2.**
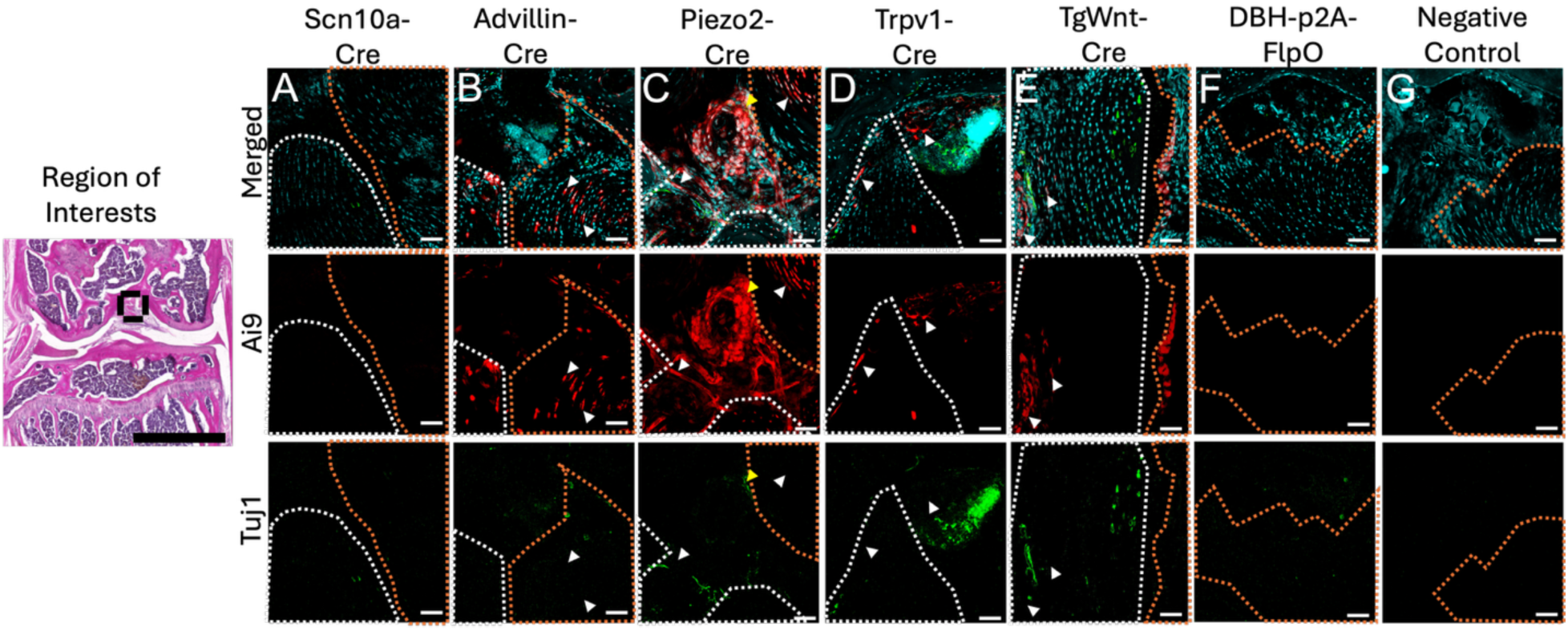
Orange dash represent ACL while white dash represents PCL. Scale bar for the images is 50 um. Yellow arrow indicates positive neuronal signal overlap between Tuj1 1 and TdTom. White arrow indicates false positive neuronal signal indicated by the lack of tuj1 and td tom overlap. Yellow arrow heads indicated reporter and NeuN colocalization. Every column represents a different genetic reporter screened in this study. Knee sections of the same area were taken from Ai9 mice (no driver) as a negative for cre recombinase and primary Tuj1 antibody (G.).

**Supplemental Figure 3.**
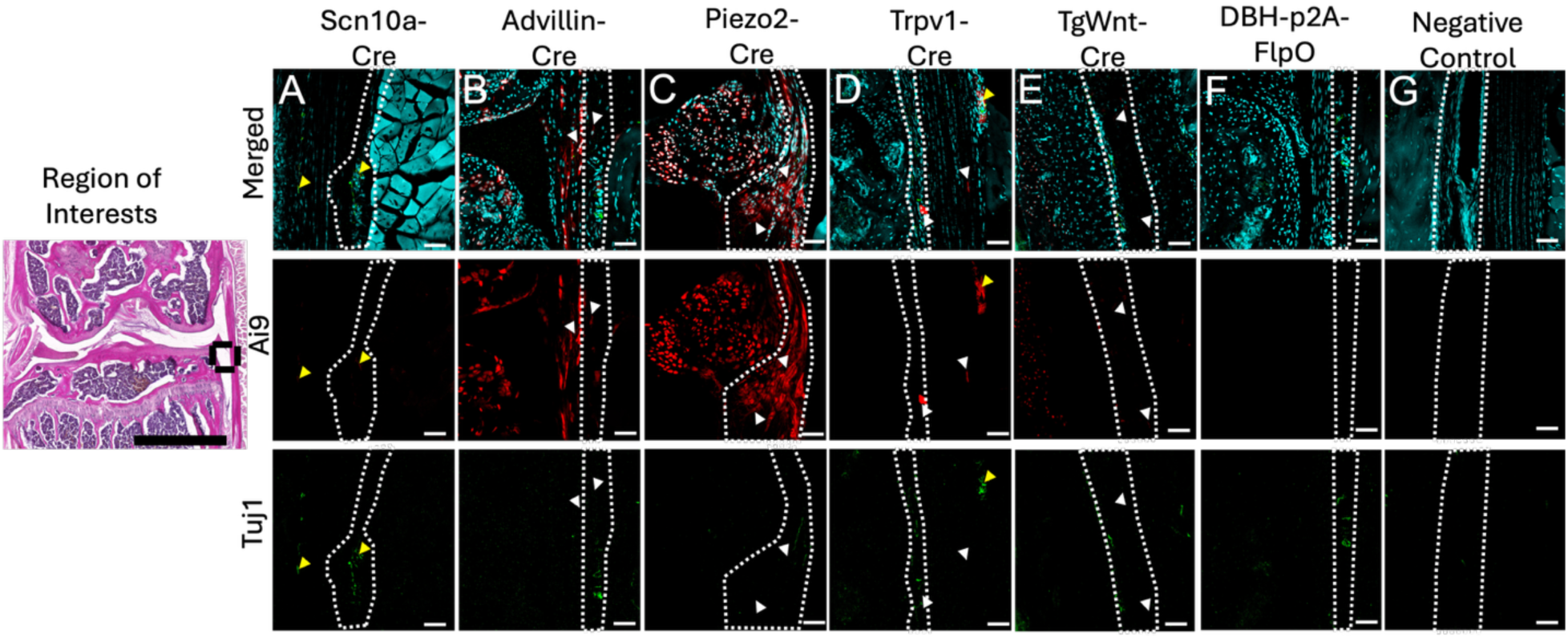
White dash represent medial synovium region. Scale bar for the images is 50 um. Yellow arrow indicates positive neuronal signal overlap between tuj1 1 and td tom. White arrow indicates false positive neuronal signal indicated by the lack of tuj1 and td tom overlap. Yellow arrow heads indicate reporter and NeuN colocalization. Every column represents a different genetic reporter screened in this study. Knee sections of the same area were taken from Ai9 mice (no driver) as a negative for cre

**Supplemental Figure 4:**
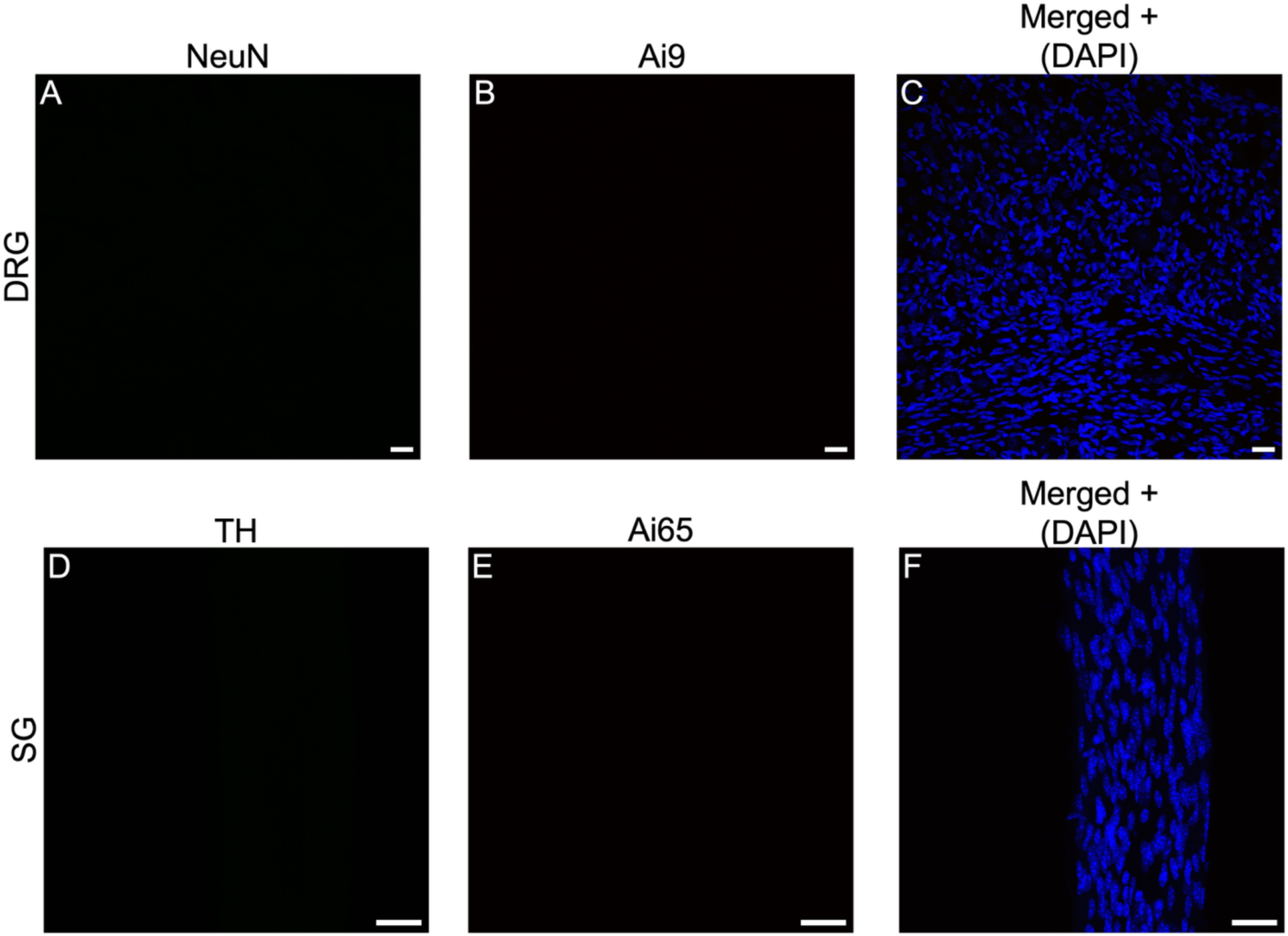
Representative images taken of DRG and SG sections from reporter strains, Ai9 and Ai65F not carrying recombinase driver alleles. Panel A represents negative control for NeuN antibody used to stain neurons in DRG sections throughout study. Image in panel B was taken with 561nm channel in Ai9 DRG section. Panel D represent negative control for Tyrosine Hydroxylase antibody stain for sympathetic neurons. Panel E shows fluorescence from Ai65F reporter strain using 561nm channel. Panels C and F represent merged channels with nucleic acid stain, DAPI. Panels D-F zoomed in 2x. Images are Z-projections spanning 20 microns. Scale bar = 25 microns.

