## Supplemental figures for "A validated set of neural gene reporter mice and chemical tracer tools for mapping knee innervating neurons"

### Slide 1
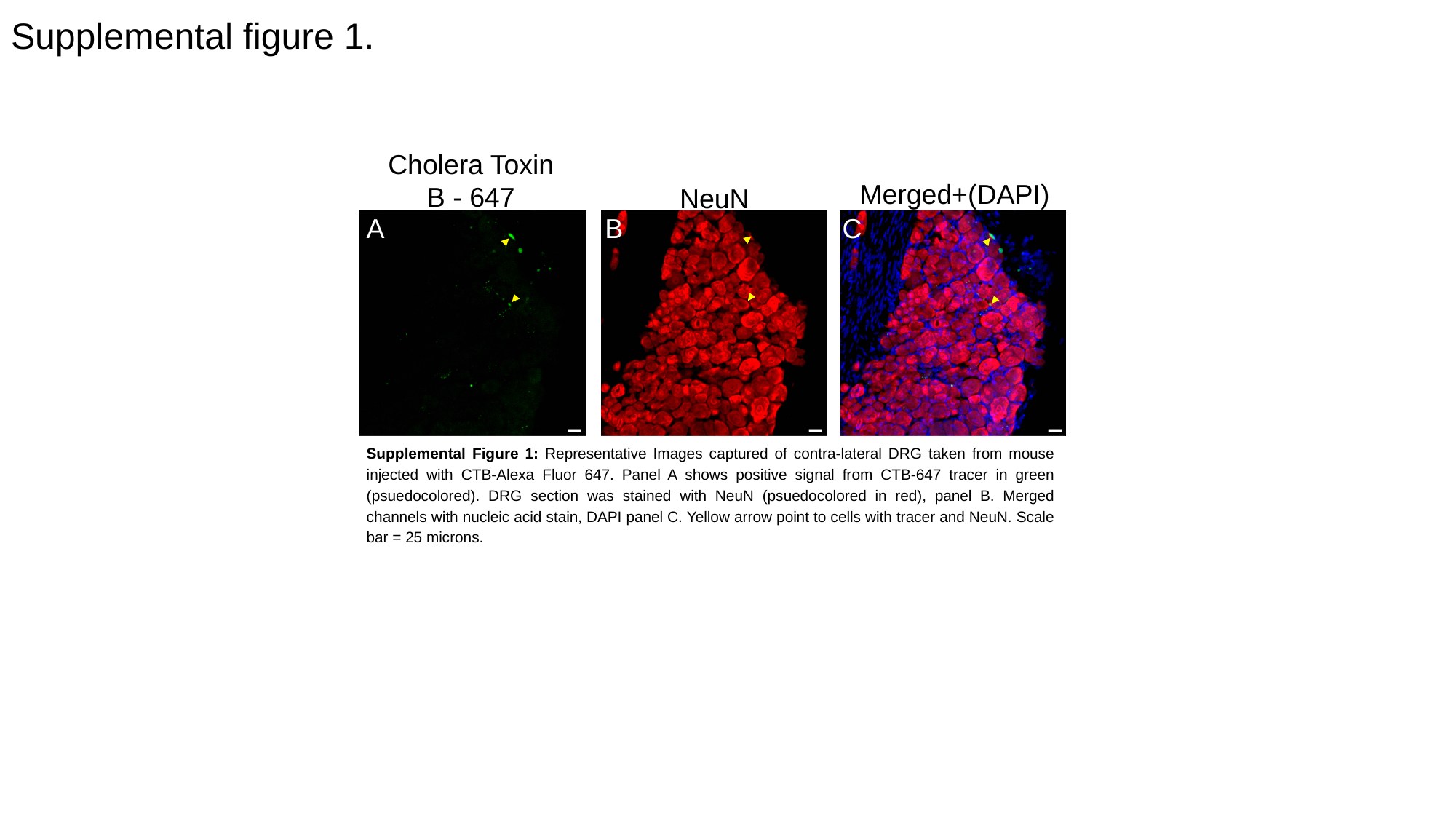

#### Supplemental figure 1.
Cholera Toxin B - 647
Merged+(DAPI)
NeuN
A
B
C
Supplemental Figure 1: Representative Images captured of contra-lateral DRG taken from mouse injected with CTB-Alexa Fluor 647. Panel A shows positive signal from CTB-647 tracer in green (psuedocolored). DRG section was stained with NeuN (psuedocolored in red), panel B. Merged channels with nucleic acid stain, DAPI panel C. Yellow arrow point to cells with tracer and NeuN. Scale bar = 25 microns.

### Slide 2
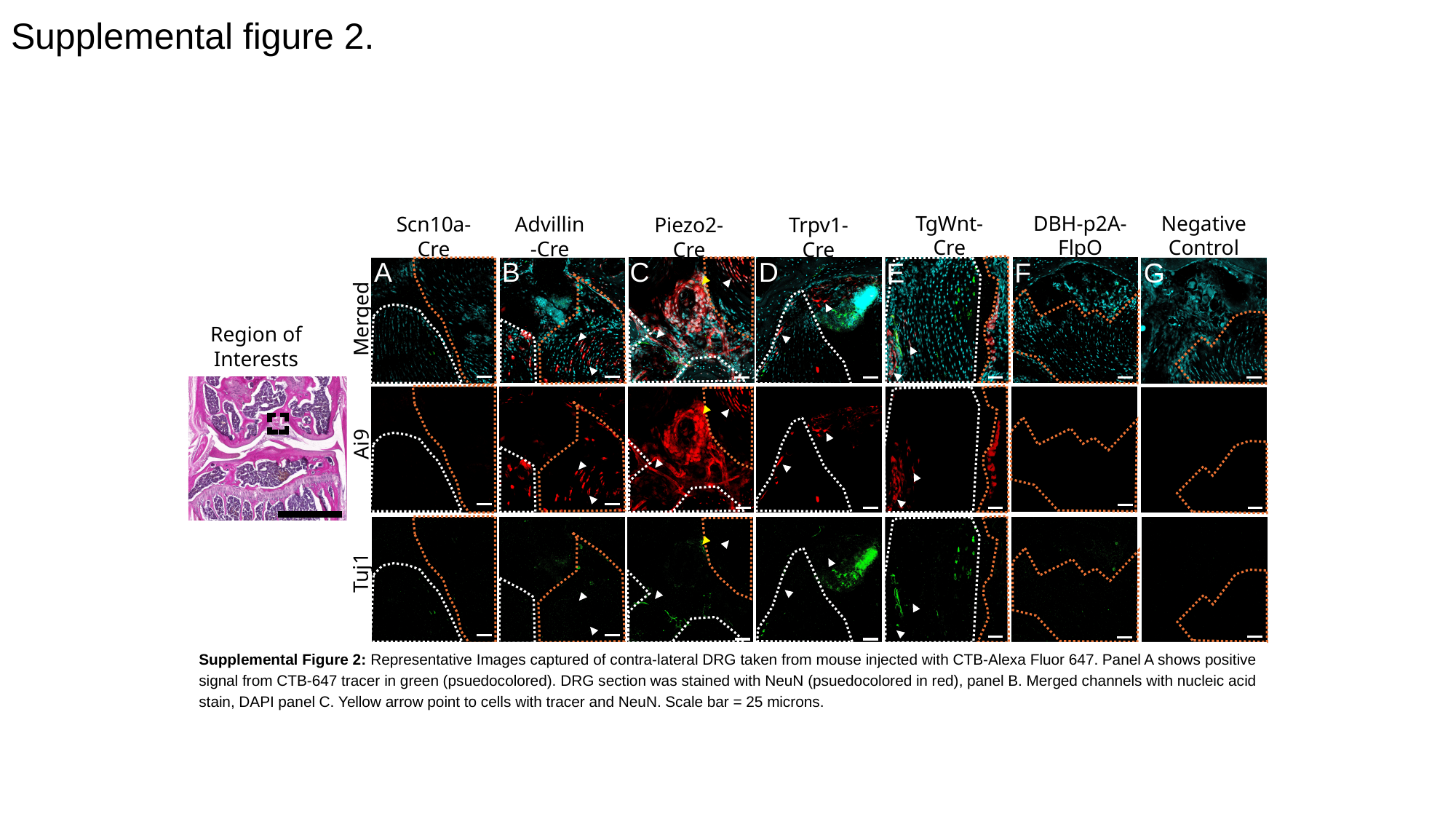

#### Supplemental figure 2.
DBH-p2A-FlpO
Negative Control
TgWnt-Cre
Advillin-Cre
Scn10a-Cre
Piezo2-Cre
Trpv1-Cre
Merged
Region of Interests
Ai9
Tuj1
A
B
C
D
E
F
G
Supplemental Figure 2: Representative Images captured of contra-lateral DRG taken from mouse injected with CTB-Alexa Fluor 647. Panel A shows positive signal from CTB-647 tracer in green (psuedocolored). DRG section was stained with NeuN (psuedocolored in red), panel B. Merged channels with nucleic acid stain, DAPI panel C. Yellow arrow point to cells with tracer and NeuN. Scale bar = 25 microns.

### Slide 3
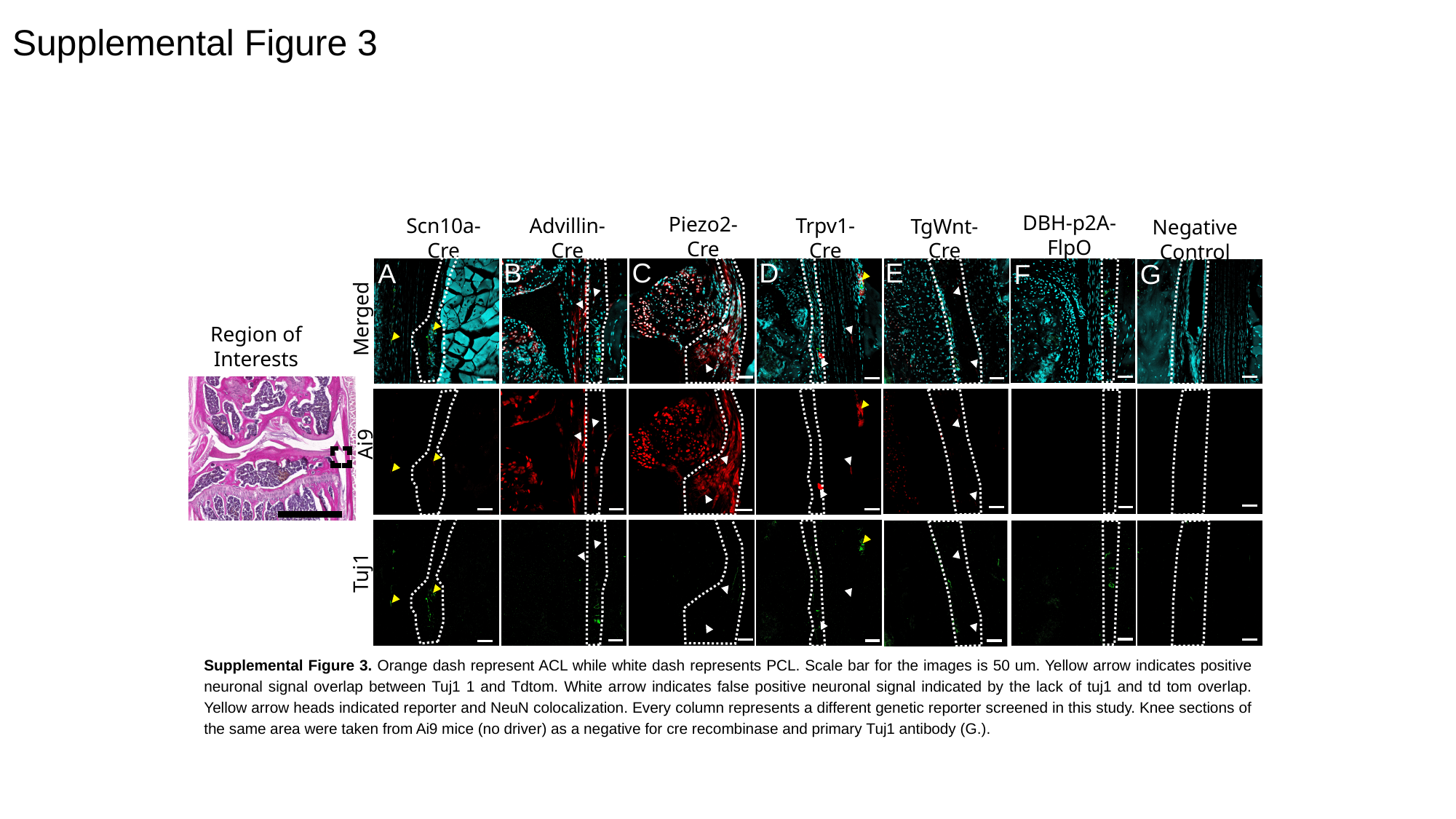

#### Supplemental Figure 3
DBH-p2A-FlpO
Piezo2-Cre
Scn10a-Cre
Merged
Ai9
Tuj1
Advillin-Cre
Trpv1-Cre
TgWnt-Cre
Negative Control
Region of Interests
C
D
E
B
A
F
G
Supplemental Figure 3. Orange dash represent ACL while white dash represents PCL. Scale bar for the images is 50 um. Yellow arrow indicates positive neuronal signal overlap between Tuj1 1 and Tdtom. White arrow indicates false positive neuronal signal indicated by the lack of tuj1 and td tom overlap. Yellow arrow heads indicated reporter and NeuN colocalization. Every column represents a different genetic reporter screened in this study. Knee sections of the same area were taken from Ai9 mice (no driver) as a negative for cre recombinase and primary Tuj1 antibody (G.).

### Slide 4
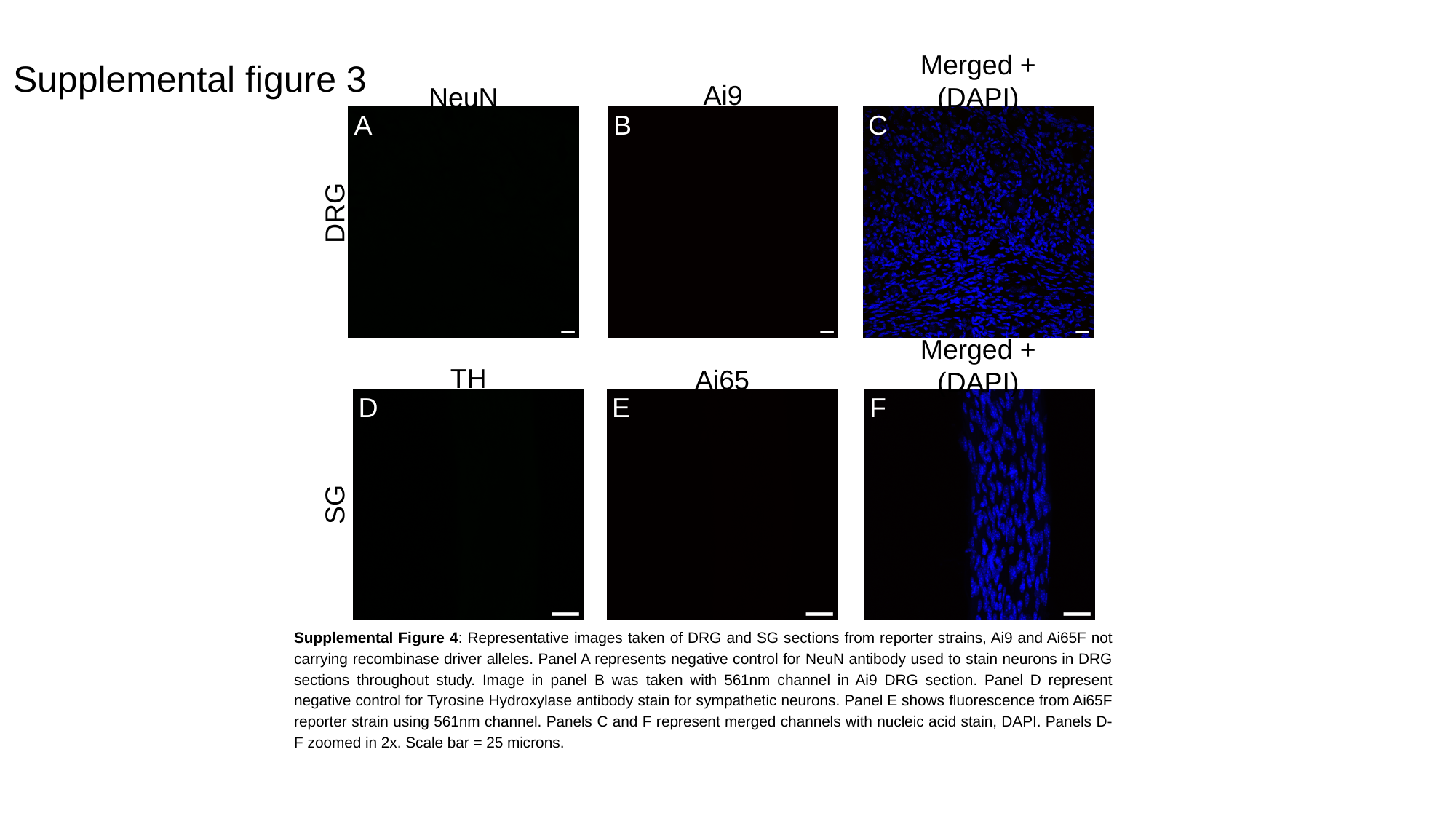

#### Supplemental figure 3
Merged + (DAPI)
Ai9
NeuN
A
B
C
DRG
Merged + (DAPI)
TH
Ai65
D
E
F
SG
Supplemental Figure 4: Representative images taken of DRG and SG sections from reporter strains, Ai9 and Ai65F not carrying recombinase driver alleles. Panel A represents negative control for NeuN antibody used to stain neurons in DRG sections throughout study. Image in panel B was taken with 561nm channel in Ai9 DRG section. Panel D represent negative control for Tyrosine Hydroxylase antibody stain for sympathetic neurons. Panel E shows fluorescence from Ai65F reporter strain using 561nm channel. Panels C and F represent merged channels with nucleic acid stain, DAPI. Panels D-F zoomed in 2x. Scale bar = 25 microns.
